# In silico chemical profiling and identification of neuromodulators from *Curcuma amada* targeting Acetylcholinesterase

**DOI:** 10.1101/2020.02.22.960732

**Authors:** Md. Chayan Ali, Yeasmin Akter Munni, Raju Das, Marium sultana, Nasrin Akter, Mahbubur Rahman, Md. Nazim Uddin, Kantu Das, Md. Hossen, Md. Abdul Hannan, Raju Dash

## Abstract

Curcuma amada or Mango ginger, a member of the Zingiberaceae family, has been revealed as a beneficiary medicinal plant having diverse pharmacological activities against a wide range of diseases. Due to having neuromodulation properties of this plant, the present study characterized the secondary metabolites of Curcuma amada for their drug-likeness properties, identified potent hits by targeting Acetylcholinesterase (AChE) and revealed neuromodulatory potentiality by network pharmacology approaches. Here in silico ADMET analysis was performed for chemical profiling, and molecular docking and molecular dynamics simulations were used to hit selection and binding characterizations. Accordingly, ADMET prediction showed that around 87.59% of compounds processed drug-likeness activity, where four compounds have been screened out by molecular docking. Guided from induced-fit docking, molecular dynamics simulations revealed phytosterol and curcumin derivatives as the most favorable AChE inhibitors with the highest binding energy, as resulted from MM-PBSA analysis. Furthermore, all of the four hits were appeared to modulate several signaling molecules and intrinsic cellular pathways in network pharmacology analysis, which are associated with neuronal growth survival, inflammation, and immune response, supporting their capacity to revert the condition of neuro-pathobiology. Together, the present in silico based characterization and system pharmacology based findings demonstrate Curcuma amada, as a great source of neuromodulating compounds, which brings about new development for complementary and alternative medicine for the prevention and treatment of neurodegenerative disorders.

## 1 Introduction

Acetylcholinesterase (AChE) is a hydrolase enzyme, which catalyzes neurotransmitter acetylcholine and some other choline ester into acetate and choline (1, 2). This catalyzing reaction terminates neuronal signals transmitted by these neurotransmitters (3, 4). Besides, AChE responses to several cellular insults like cellular stresses and also disrupts cell proliferation and differentiation (3). The active site of AChE is located in the deep and narrow gorge side of this enzyme, which consists of anionic and esteratic subsite acts as catalytic and choline-binding site respectively (5, 6). The substrate binds to the hydrophilic and uncharged anionic subsite (3). The positively charged substrates have no interactions with the anionic negatively charged amino acid residues, however, interact with the anionic active site, containing 14 amino acids. Among these amino acids, TRP^84^ plays a vital role in substrate binding (6). Since the AChE impedes the normal signaling system, its interference is necessary in order to maintain the ACh’s persistence action. Due to its higher catalytic activity (25,000 reactions per seconds), AChE received considerable attention from researchers (3). In recent decades, treatments for neurological disorders have been designed to treat cholinergic dysfunction primarily based on AChE (AChE-I) inhibitors, which improve cholinergic efficiency and have persistent therapeutic effects (7). A recent study suggest that AChE inhibitors up regulate cholinergic function which was associated with the suppression of proinflammatory cytokine and lymphocyte proliferation (8). Several AChE inhibitors are available in the market, such as biperiden, donepezil, edrophonium, galantamine, huperazine, neostigmine, rivastigmine, tacrine, and trichlorfon (1, 9–12). These drugs are reported to improve cognitive function and approved for different neurodegenerative disease (NDDs) therapy such as Alzheimer’s Disease (AD), Parkinson’s Disease (PD), Multiple Sclerosis (PD), Huntington’s disease (HD) and prion diseases (4, 13–15). However, some of these drugs shows various side effects including, hepatotoxicity (tacrine) (16), anorexia, abdominal pain, diarrhea, gastrointestinal anomalies-nausea and bradycardia (donepezil) (17). Rivastigmine and galantamine causes anorexia, diarrhea, dizziness, headache, nausea, vomiting, abdominal pain, and syncope (18, 19), whereas biperiden causes bowels and bladder atonic states, confusion, agitation, mydriasis, red face, hyperthermia, dryness of mucous membranes, and hallucinations (20). Thus there is a need for effective and safe AChE inhibitor, which can prevent neurodegeneration, block the disease progression at the primary stage or improving the cognitive impairment conditions to reduce the social and economic cost of caring NDDs patients and improving the quality of life.

Natural products derived from resources have been proved as a very efficient therapeutic agents due to their antioxidant, antiapoptotic, antiamyloidogenic, and mitochondrial function modulatory effects (21, 22) and some recent findings suggest that they have counter action on protein misfolding, also influence proteasomal breakdown, aggregates clearance and autophagy (23, 24). Plants provide a great source of alkaloidal, polyphenolic and terpenoid compounds with better therapeutic potentials likewise, anxiolytic and relaxant properties of *Lippia citriodora* derived compounds (25), nitric oxide and inflammation inhibition by *Chionanthus retusus* extracts (26), and apoptosis inhibitory effect of *Centella asiatica* extracts (27). *Curcuma amada* Roxb., from the family of Zingiberaceae, was reported to show numerous pharmacological activity including antibacterial, antifungal, analgesic, anti-inflammatory, anticancer, antihyperglyceridemic, free radical scavenging, and also superoxide scavenging activities (28–30).

This plant is similar to *Curcuma longa* and has raw mango-like aroma in rhizome and thus they also called mango ginger (31). The recently published report showed that the ethanol extract from the rhizome processed the depressant activity in CNS and potential antinociceptive activity (32). Besides, the hydroalcoholic extract was responsible for the neuroprotective activity (33), which ultimately indicates the presence of various classes of secondary metabolites such as steroidal lactones, phytosterols, sitoindosides and alkaloid in *C. amada* that may modulate the cognitive and memory function in brain by modulating expressions of cholinergic and glutamatergic systems. Therefore, the present study aimed to explore the acetylcholinesterase inhibitors from the various reported secondary metabolites of *C. amada* to prevent the development of cognitive impairment, improve the physiological condition of NDD patients.

## 2 Materials and Methods

### 2.1 Collection and preparation of ligands

The information regarding the secondary metabolites of *Curcuma amada* was assembled from literature and web resources in order to develop an exclusive curated chemical library. In this regard, literature mining was done against international databases, including Google Scholar, Scopus, Web of Science, and PubMed. The obtained information was dual checked, corrected and respective chemical structures were obtained from available chemical databases, including ‘Pubchem’ and ChemSpider’. The chemical name, IUPAC name, plant part and structural information were also collected for the selected compounds. The structures, which are not available in the chemical databases, were drawn by the Chemdraw software. All structures were converted to three dimensional by using Ligand preparation wizard of Maestro 11.1 (34) with an OPLS 3 force field. Their ionization states were generated at pH 7.0±2.0 using Epik 2.2 in Schrödinger Suite (Schrödinger, LLC, New York, NY, USA) (35).

### 2.2 Determination of ADMET properties

The QikProp program (QikProp, Schrödinger, LLC, New York, NY, USA) was used to calculate ADMET properties (36). Total of 48 molecular descriptors were calculated using this program in a normal mode. At first, QikProp generates physical descriptors and later calculate ADMET. The pharmacokinetic profiles of the compounds are determined by using an all-round ADME compliance parameter (indicated by # stars). The #stars parameter shows the number of property descriptors that QikProp estimates to fall outside the optimum range for 95% of recognized medicines (37).

### 2.3 Virtual Screening

Virtual screening was done by the molecular docking approach through AutodockVina software. Initially, the structure of AChE was retrieved from the RCSB protein databank (https://www.rcsb.org, PDB ID: 4EY7) and prepared by cleaning water molecules, ligands and energy minimizing by steepest descent and conjugate gradient techniques. GROMACS 96 43B1 parameters were used for all of these calculations in the *in-vacuo* system by the SWISS-PDB viewer (38). All 3D models were produced and minimized using UCSF Chimera software (Amber Force field) in the charged form (39). In the Vina docking procedure, the number of binding modes and Exhaustiveness were set to 100 and 25 respectively (40) The AutoDock tools from MGL software suite have been used to convert pdb file into pdbqt. The grid box size was kept at 58.8163, 61.2067, and 72.8274 respectively for X, Y, Z. axis in AutoDockVina and. Docking calculations were set to 50 runs. A population of 150 individuals and 2,500,000 function evaluations were used. By using a genetic algorithm, the structure optimization was done for 27,000 generations. Maximum root mean square tolerance for conformational cluster analysis was 2.0 Å. AutoDockVina performed cluster analysis at the end of calculation (41). AutoDockVina developers provide shell script implementing through AutoDockVina. The results were evaluated binding energy values by sorting Kcal/Mol as a unit for a negative score of different ligand-protein complexes.

### 2.4 Induced Fit Docking analysis

To exceed the docking analysis with more accurate bioactive conformation, the Schrodinger’s Induced Fit Docking (IFD) was applied. After the virtual screening, IFD docking was carried out by using the IFD module of Schrödinger-Maestro v11.1 (42). By minimizing the receptor with an RMSD comprising limit 0.18 Å, the glide docking was run and a box size was generated automatically. For both the ligands, the van der Waals scaling factors were set to 0.50 and 0.70, respectively to soften the potential of both ligands and receptors and the B-factor side chains were trimmed automatically. The number of maximum of 20 poses were kept during the docking simulation. To bind the domain flexibility, a cutoff of 5 Å was set for all residues of ligand poses which were refined using Prime molecular dynamics module and the side chains were again optimized. The best poses within 30.0 kcal/mol were then re-docked generating overall 20 structures with the protein molecule by using the glide SP. Finally, for each ligand induced-fit receptor docking generate an IFD score in which the lowest IFD score containing poses were then selected for further study.

### 2.5 Molecular Dynamics Simulations (MDS)

Molecular dynamics simulation was run for the protein-ligand complexes derived from IFD, using YASARA dynamics software. The simulation protocol began with the hydrogen-bond network optimization, and subsequently, a cubic simulation cell in the periodic boundary condition was generated, where the protein of each complex was parameterized by the AMBER14 force field (43, 44). The ligand was parameterized utilizing AutoSMILES (45) algorithm, which automatically creates topology of unknown organic molecules through semi-empirical AM1 calculation of mulliken point charges (45) in COSMO solvation model, assignment of atom and bond types to AM1BCC (46) and also assignation of GAFF (General AMBER Force Field) (47). Using the TIP3P water model, the simulation box was solvated by maintaining a density of 0.997 g/L. During solvation, the pH of the system was maintained at 7.4 to mimic the physiological conditions, and accordingly, the protonation states of each amino acid residue were determined in a combination of the H-bonding network and SCWRL algorithm, which employs dead-end elimination and graph theory (48). Furthermore, the solvation system was supplemented with Na^+^ and Cl^−^ ions (49). In order to eliminate conformational stress, the energy minimization protocol used to minimize the initial structure through steepest descent without electrostatic interaction. After this, a steep descent minimization was incorporated to relax the structure, which was subject to the total potential reduction of energy over 5,000 cycles, until convergence was achieved. The energy per atom was enhanced in 5000 steps by less than 0.05 kJ / mol. In order to describe long-range electrostatic interactions at a threshold distance of 8Å in physiological conditions (298 K, pH 7.4, 0.9 % NaCl), MD simulations were carried out using PME methods (50). The simulation time step interval was set to 2.0 fs together with multiple time step algorithm (51). MDS were done for 50 ns and the trajectories were saved in each 50 ps with a constant pressure and Berendsen thermostat. The subsequent trajectories were analyzed for consistency with various analytical steps, including RMSD (Root Mean Square Deviation), RMSF (Root Mean Square Fluctuation) using YASARA developed into macros, VMD (52) and Bio3D (53) software. After that, the MM-PBSA binding free energy calculations were done by in built macro of YASARA dynamics software, where the resulted binding energy is calculated through following equation,

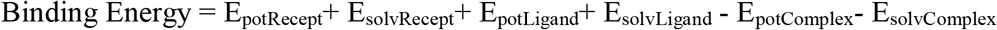

### 2.6 Free Energy Landscape (FEL)

Free Energy Landscape (FEL) is a mapping system of all possible conformations of molecules in a system which is used to bring out their corresponding energy levels especially Gibbs free energy. In this study, the protein stability is expressed through Gibbs Free Energy by determining the function of protein enthalpy and entropy (54, 55). FEL analysis was also carried out to check the evaluation of the trajectory changes. In this study, the FEL was investigated by following equation:

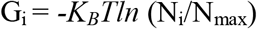

Where, K_B_ is Boltzmann’s constant, T is temperature (300K), *N*_*i*_ is the population of bin *i* and *N*_max_ is the population of the most populated bin. Color code modes were utilized to discuss different energy levels.

## 3 Results

### 3.1 Database and In silico Chemical Profiling

In our study, we curated 137 compounds from different parts of the *C. amada* that are extracted and reported in previous studies. The details of these compounds are depicted in the supplementary **file 1** with their chemical name, IUPAC name, and 2D structure. These compounds contain 81% essential oils, 7% phenol and terpene, 4% terpenoid and 2% curcuminoid (Figure 1a). The physical and chemical profiling of these compounds were analyzed, where the result revealed that the molecular weight of our isolated compounds was in the acceptable range (100%) (Figure 1b) whereas, 86.13% compounds (total 118 individual compounds) molecular weight was in this interval 100-200 (Figure 1b). According to ‘rule of five’ postulated by Lipinski, Number of Hydrogen Bond Acceptor (HBA) and Number of Hydrogen Bond Donor (HBD) should not be above 10 and 5 respectively. The HBA (Figure 1c) and HBD (Figure 1d) depicted that, all of our isolated compounds showed the acceptable range of HBA and HBD, moreover, 37.23% and 62.77% compounds showed a peak value of 0 for both HBA and HBD. As higher polar compounds cannot pass blood brain barrier (BBB), therefore, the prediction of BBB penetration is necessary. Polarity based accession to CNS and BBB permeability of the compounds were indicated using log B/B (reference range −3 to 1.2) (56, 57).

**Figure 1:**
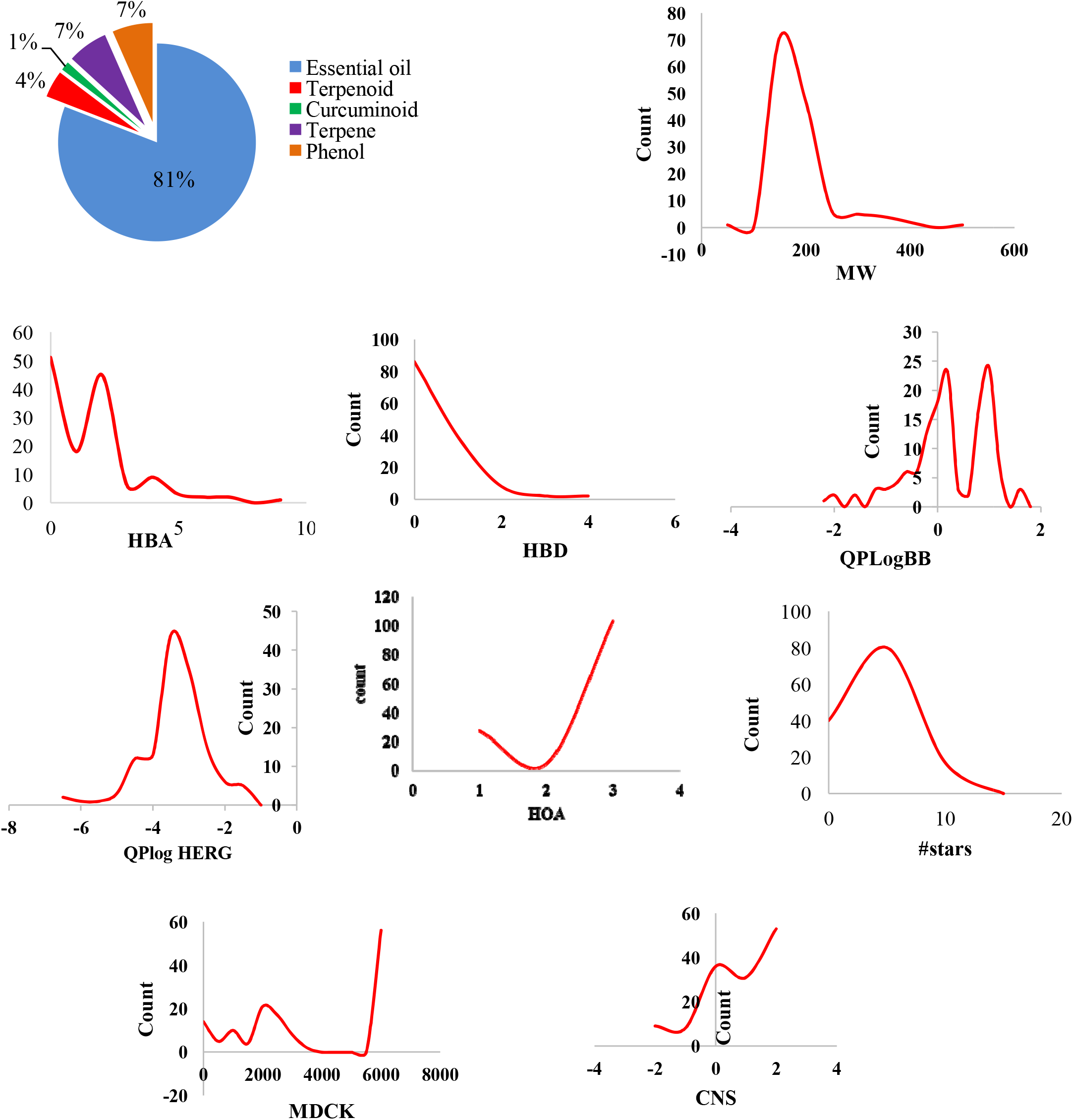
(a) Pi chart showing the classification of *Curcuma amada* compounds. Distribution curves of (b) Molecular weight against count, (c) Hydrogen bond acceptor against count, (d) Hydrogen bond donor against count, (e) Calculated QP log BB against count (f) Plot of predicted logHERG values against count, (g) Calculated percentage of Human oral absorption against count, (h) #stars against count, (i) Plot of logMDCK values against count, (j) Plot of CNS against count

The QikProp analysis depicted that about 92.07% compounds have the ability to cross the BBB, while 87.59% compounds showed the ability to activity in CNS system (Figure 1e and 1j). The cardiac toxicity assessment is also necessary for a drug-likeness substance. Human Ether-a-go-go related gene (hERG) gene is responsible for maintaining the cardiac systolic and diastolic activity by maintaining the potassium ion channel. In this study, we estimated IC50 values to model in silico toxicity of the drug-likeness compounds for blocking the HERG K+ channel. About 94.89% of compounds were within the recommended range (logHERG > −5) (Figure 1f). The compound’s bioavailability relies on the compounds absorption processes and their metabolism (56). The oral absorption assessment was predicted by calculating Human Oral Absorption (HOA). It was predicted that 75.91% of compounds have higher HOA (attributed to a value of 3), 3.65% compounds have medium HOA (attributed value is 2). The rest 20.44% of compounds have low absorption (Figure 1g). Besides, to estimate oral abortion Madin-Darby canine kidney (MDCK) cells are widely used because they can express transporter proteins with low level of enzymes expression. Evaluation of MDCK cell permeability is an additional criteria for analyzing BBB penetration. About 86.13% isolated compounds have MDCK cell permeability (Figure 1i).

The overall drug-likeness properties of *C. amada* compounds have been evaluated by #stars parameter. Distribution graph for this parameter shown in (Figure 1h). This graph depicted that overall 87.59% compounds have drug-likeness properties.

### 3.2 Virtual Screening

In the area of in silico drug design, virtual screening is a vital approach to screen out the notable hits from thousands of compounds (58, 59). Virtual screening is proposed to be a successful alternative methods of lead compound identification in drug discovery process (60, 61). It has received considerable interest in lead detection, since it looks like biological research using computer technology that appear to be closer to true value (62). In this case, the docking protocol describes the ligand-binding site in the receptor and also the ligands specification in the particular location.

All identified (137) compounds were imported for docking calculation where we found four compounds with lowest binding energy. These four compounds were selected for further analysis. According to docking score graph of isolated compounds from *C. amada*, more than 45 compounds showed docking score of −6 while, 40 compounds showed the docking score of −7. Again, there were 30 and 8 compounds for the docking score of −8 and −9 respectively (Figure 2a). Among the four selected compounds, the highest docking score was found −14.523kcal/mol for curcumin, and demethoxy curcumin, bisdemethoxycurcumin, and β-sitosterol were scored −12.993 kcal/mol, −11.913 kcal/mol, and −10.729 kcal/mol respectively (Table 1). Therefore, Curcumin and demethoxy curcumin showed higher binding affinity while bisdemethoxycurcumin and β-sitosterol showed lower binding affinity towards AChE. These compounds are considered for further analysis

**Figure 2:**
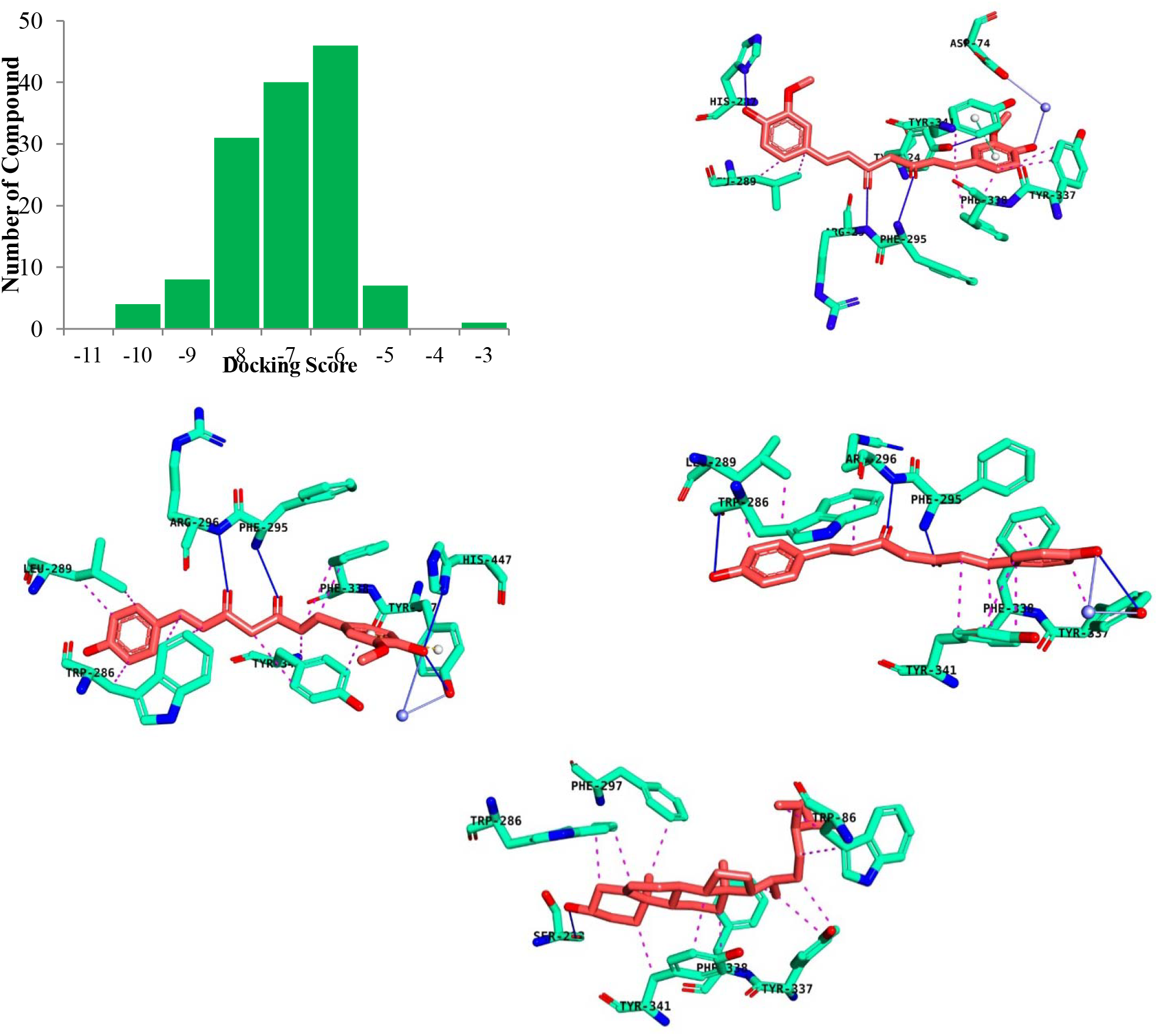
(a) Docking score of selected compounds. Molecular interactions between AChE and selected compounds. (a) AChE-Curcumin, (b) AChE-Demethoxycurcumin, (c) AChE-Bisdemethoxycurcumin, and (d) AChE-β-Sitosterol

### 3.3 Binding interaction analysis by IFD study

In docking calculations, the rigid receptor does not always provide the exact binding pattern of ligand on the active site. Because in reality, the protein undergoes some spatial changes. This mobility can be due to the flexibility of the active site residues, which is also complementary to their side chain. This can be considered in induced-fit process. IFD is a combination of both molecular dynamics and molecular docking, predicts accurately receptor natural conformation changes and ligand binding modes. Accordingly, the top 4 hits were allowed to the induced fit docking. Ligand pose having the lowest IFD score were selected for the molecular interaction analysis. The IFD docking scores are represented in **Table S1**.

Figure 2b represents the molecular interactions between the curcumin and AChE, in which the oxygen atom of the 3-methoxy group at phenyl ring formed hydrogen bond with TYR^124^ at a corresponding distance of 2.32112 Å. This ring also formed pi-pi stacked bond with TYR^341^. The two oxygen atoms of curcumin showed three hydrogen bonds with PHE^295^ and ARG^296^ residues with the bond lengths of 2.11364Å, 2.59759Å, 1.88951Å, respectively, here PHE^295^ involved in duel hydrogen bonding. In contrast, two hydrogen bonds were observed with HIS^287^ and pi-alkyl bond with LEU^289^ at 7^th^ (4-hydroxyl-3-methoxyphenyl) ring. The pi-alkyl bond also formed between curcumin and PHE^338^, TYR ^337^ residues.

In demethoxy curcumin, PHE^295^, ARG^296^ hydrogen bond were seen in (Figure 2c) with the part of 1, 6-diene-3, 5-Dione whereas, PHE^295^ containing two hydrogen bonds observed with the distance of 2.03758Å, 2.48401Å. On the contrary, hydroxyl group formed hydrogen bond with HIS^447^ and pi-alkyl bond formed with PHE^338^ and TYR^337^ at 3-methoxy group. Others residue, TRP^286^ was involved in pi-alkyl bond and LEU^289^ was involved in alkyl-alkyl bond at phenyl ring.

In case of bisdemethoxycurcumin (Figure 2d), hydrogen bonds were observed at 1, 6-diene-3, 5-Dione with PHE^295^ and ARG^296^ residues, while PHE^295^ formed two hydrogen bonds with the distance of 2.03758Å and 2.48401Å. Hydrophobic interactions were seen with TYR^337^, TYR^341^, PHE^338^, and TYR ^337^at the phenyl ring. Additionally, TRP^286^ formed hydrogen bond with 4 hydroxyl group and pi-alkyl bond with aromatic ring. Besides alkyl-alkyl bond was associated with LEU^289^.

In beta-Sitosterol major hydrophobic interactions were observed with TRP^86^, TRP^286^, PHE^297^, TYR^337^, PHE^338^, and TYR^341^ residues by means of pi-alkyl bonding (Figure 2e). A single hydrogen bond was observed in cyclophenanthren-3-ol with the residue SER^293^ having the bond length of 2.18947Å.

The catalytic active site (CAS) is built by several amino acid residues including tryptophan, phenylalanine, glutamate, histidine, and serine. A hydrophobic region, the peripheral anionic site (PAS) traps ligands and sends them to the deep cat lytic region. Several studies have been conducted in noncompetitive AChE inhibition and the role of amino acid residues during this process (63–65). The inhibition requires binding of inhibitors to the active site george (63, 66). Some previous study related to AChE inhibition depicted that inhibitors interact with the following residues Trp^86^ and Phe^338^ of CAS and residues Trp^286^, PHE^295^, Tyr^337^, and Tyr^341^ of PAS (67–69). From the docking analysis, it can be concluded that all the studied compounds formed hydrogen and hydrophobic interactions with TYR^337^, TYR^341^ and Phe^338^ residues which are involved either in the peripheral site or in the ligand recognition mechanism by allosteric activation. Our selected compounds also interacted with the previously studied active site residues. Thus, they can successfully block the active-site gorge of the AchE. To validate and reestablish the result of docking analysis and to see whether the interacted residues will be fully conserved or not, MDS was performed.

### 3.4 Molecular Dynamics Simulation

To evaluate the motion and trajectory of molecules, structural features and conformational change of molecules, we conducted MD simulation for 50 ns. In this study, four protein-ligand complexes of beta-sitosterol, bisdemethoxy carcumin, demethoxy curcumin and curcumin were subjected to MD simulation by means of RMSD analysis of the Cα-atoms, critical residues fluctuations analysis by RMSF, number of hydrogen bond and MM-PBSA for the analysis of binding free energies.

#### 3.4.1 Analysis of protein’s stability and compactness

Protein’s overall stability and conformational difference after binding of the studied compounds was evaluated by RMSD analysis of the Cα-atoms of the respective protein (70). The protein will be stable if the deviation is shorter. According to the result, upon binding of four compounds, the stability of the AChE ranges between 0 to 2 Å. The average deviation for AChE-β-sitosterol, AChE-Bisdemethoxycurcumin, AChE-Demethoxycurcumin and AChE-Curcumin were observed to be 1.4, 1.25, 1.2 and 1.1 Å respectively, though the AChE-Curcumin complex showed more fluctuations throughout the simulation (Figure 3a). Thus the AChE-Curcumin complex showed lower RMSD with more fluctuations than the other three complexes indicating more significant binding. Again, all ligands appeared the same flexibilities during the simulation though the beta-sitosterol showed the highest deviation among the four ligands which became more flexible after 26 ns. The average RMSD values for Bisdemethoxycurcumin, De-methoxycurcumin and Curcumin ligands were 0.7, 0.5 and 0.6 Å respectively and there were not many fluctuations after 26 ns for these (Figure 3b). Therefore, the overall result of ligand RMSD explains that despite the De-methoxycurcumin appeared the lower deviation from Curcumin, AChE was more stable when combining with curcumin rather than De-methoxycurcumin.

**Figure 3:**
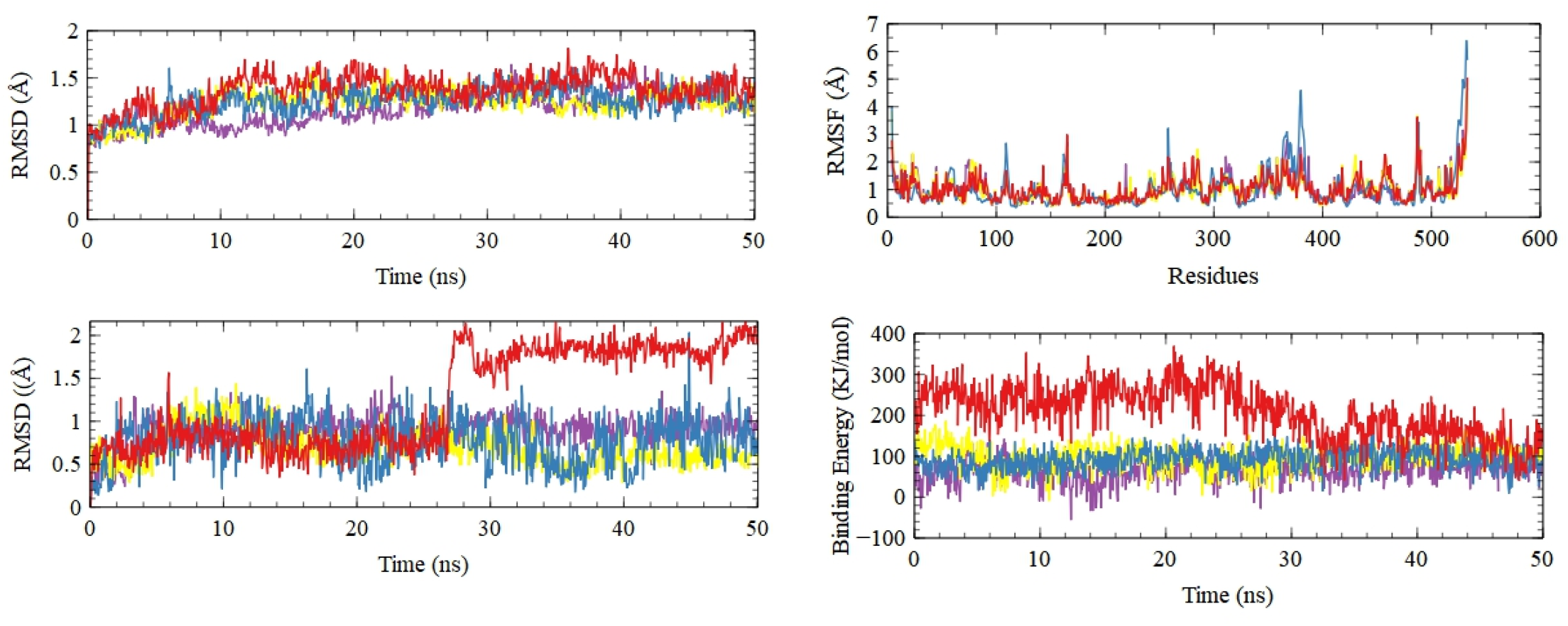
Root Mean Square Deviation (a) Protein RMSD, (b) Ligand RMSD, (c) Root mean square fluctuation (RMSF). (d) Binding Free Energy graph where, in all cases Red, Blue, Yellow and Purple color denote Beta-sitosterol, Bisdemethoxycurcumin, Demethoxycurcumin and Curcumin respectively.

The local fluctuations as well as differences in residue movements could be estimated by Root mean square fluctuations (RMSF). According to Figure 3c, AChE-Bisdemethoxy curcumin showed the highest RMSF values with the highest peak at 6.5 Å and the residues responsible for these increasing fluctuations are TYR-105, Phe-338 and VAL-379. AChE-Beta sitosterol, AChE-Demethoxycurcumin and AChE-Curcumin showed minimum RMSF, where the highest peaks were at 5.2 Å between 538-540 residues and at 4.8 Å between 525 to527 residues and 4.6 Å between 534-536 residues than AChE-Bisdemethoxy curcumin. Moreover, beta-sitosterol decreased flexibility of AChE in 86, 285-29, 325, 400 and 475-476 residues where TRP-286 residue belongs to PAS. However, fluctuations were seen to reduce for Curcumin when binding to AChE with residues with residues 95-105, 200-205, 290-298, 323-325, 445-450 and 470-475, where the residue PHE-295 and ARG-296 also made hydrogen bond with AChE. The docking study of protein-ligand complex predicted hydrogen and hydrophobic interaction with TRP-286 and GLU-293, as it was indicated in RMSF plot for lower fluctuations in AChE-Curcumin complex in comparison with other three protein-ligand complexes. The result reveals that the interaction between AChE and Bisdemethoxycurcumin is weak while the interaction between AChE and Curcumin is strong.

#### 3.4.2 Ligand binding insights

Hydrogen bonds are important mediator of ligand binding to the protein with high affinity and specificity (71). Therefore, this study also investigated hydrogen bond profiling of the studied complexes throughout the simulation and depicted in Figure 4a. From the figure 4a it was observed that, the four complexes exhibited around similar hydrogen bonding ranging from 0 to 3. AchE-curcumin displayed 0 to 2 while AchE-demethoxycurcumin displayed 0 to 1 hydrogen bonds. Similarly, AchE-beta-sitosterol showed numbers of 0 to 1 hydrogen bonds and AchE-bisdemethoxy curcumin possessed the hydrogen bond value of 0 to 2 but less denser than AChE-Curcumin complex. So, all the complexes are very much consistent with each other in case of hydrogen bonds. It can be concluded that AChE-Curcumin showed maximum hydrogen bonds, while beta-sitosterol showed minimum hydrogen bonding during the simulation and after 26 ns it didn’t show any hydrogen bond which showed several contact again after 45 ns with AChE. Furthermore, Figure 4b was plotted to show hydrogen bond occupancy where the residues are shown which were behaved as a donor or acceptor. According to figure 6, AchE-beta-sitosterol made H-bonds with several residues including TRP 286, GLU 292 and in which total H-bond occupancy was more than 5% while, AchE-bisdemethoxy curcumin formed H-bonds with residues of PHE 295, ARG 296 and showed H-bond occupancy of more than 15% throughout the whole simulation. In addition, AChE-demethoxy curcumin formed H-bond with PRO-296, ARG 296 but AChE-Curcumin formed H-bonds with HIS 447, PHE 295, ARG 296 residues where H-bond occupancy for PHE 295 was less than 40%. However, AChE-Curcumin showed maximum H-bond occupancy with several residues in comparison with other three complexes.

**Figure 4:**
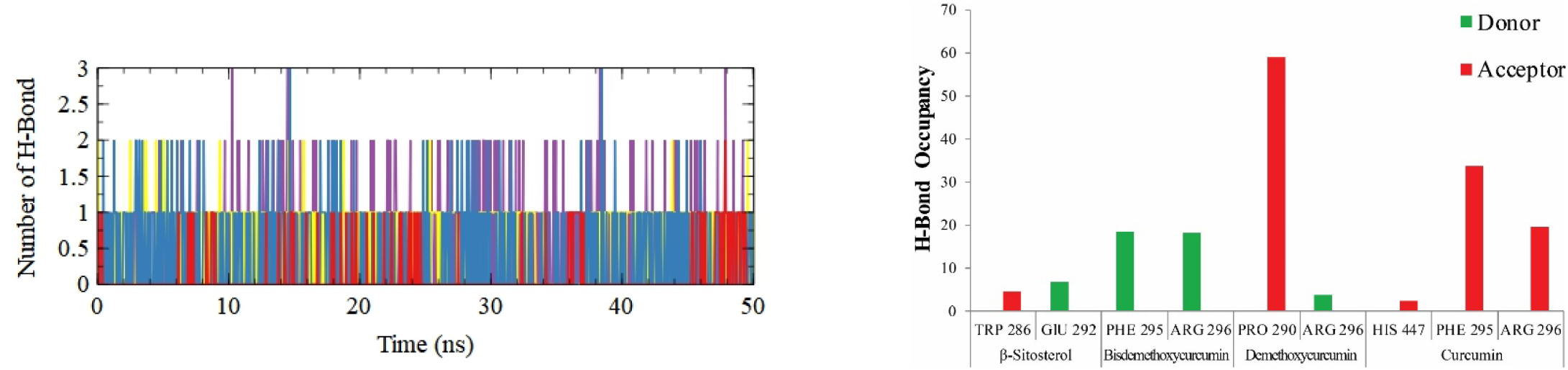
(a) Number of hydrogen bond, (b) Hydrogen bond occupancy analysis, where in all cases Red, Blue, Yellow and Purple color denote Beta-sitosterol, Bisdemethoxycurcumin, Demethoxycurcumin and Curcumin respectively.

The binding energy was also calculated to see how ligands affect the structure of AChE. The results are plotted in Figure 3d. On average, the binding energy of AChE with beta-sitosterol was 240 KJ/mol which was started to decline and after 20 ns it showed stability with binding energy of 150 KJ/mol. Again, AChE with bisdemethoxycurcumin, demethoxycurcumin and curcumin showed similar binding energy during the simulation. On average the binding energy were 80 KJ/mol, 100 KJ/mol and 50 KJ/mol, respectively. As more positive energy represents better binding, beta-sitosterol showed stronger bond with AChE. These compounds are not only stable in silico but also showed positive effect on different types of animal model system depicted in Supplementary table 2.

Overall, MD simulation study was carried out to execute the binding mechanism and dynamic stability of four compounds interacting with AChE. The whole analysis depicts that Curcumin is the best inhibitor for AChE among the four ligands because of lower RMSD value. Curcumin also showed lower RMSF due to greater hydrogen bonding with PHE 295 residue which shows also hydrophobic interaction with curcumin. Moreover, beta-sitosterol showed pi-alkyl interaction with TRP 286 residue of peripheral anionic site and TRP 286 residue also showed highest binding energy which makes it stronger and caused little configuration changes of protein by several fluctuations during the whole MD simulation. However, proteins are flexible in nature and this character accomplishes ligand binding, ligand recognition or interaction. So it can be concluded that, along with curcumin, with more flexibilities and highest binding energy, beta-sitosterol could be a better inhibitor of Acetylcholinesterase.

To interpret the conformational states related with different free energy states, the FEL was plotted by using Rg and RMSD to get the probability distribution P in Figure 5. The conformations with minimal free energy are found in the blue area which is more stable than the conformations with high free energy which is found in red area. Here, we made use of Rg and RMSD to get the probability distribution P. According to the figure, there was maximum number of basins in AChE-β-sitosterol complex with the highest energy conformational state while, The AChE-bisdemethoxycurcumin showed minimum energy basins among the four complexes with lowest energy conformational state having the most stable conformation. Furthermore, in curcumin, the presence of basins were bigger than demethoxy-curcumin at Rg of 2.262~2.278 Å with RMSD of 2.05~2.085 Å and at Rg of 2.269~2.2857 Å with RMSD of 2.06~2.095 Å respectively. Therefore, the FEL highlights the structural stability of bisdemethoxycurcumin and curcumin in complexes with AChE more than demethoxy-curcumin and β-sitosterol.

**Figure 5:**
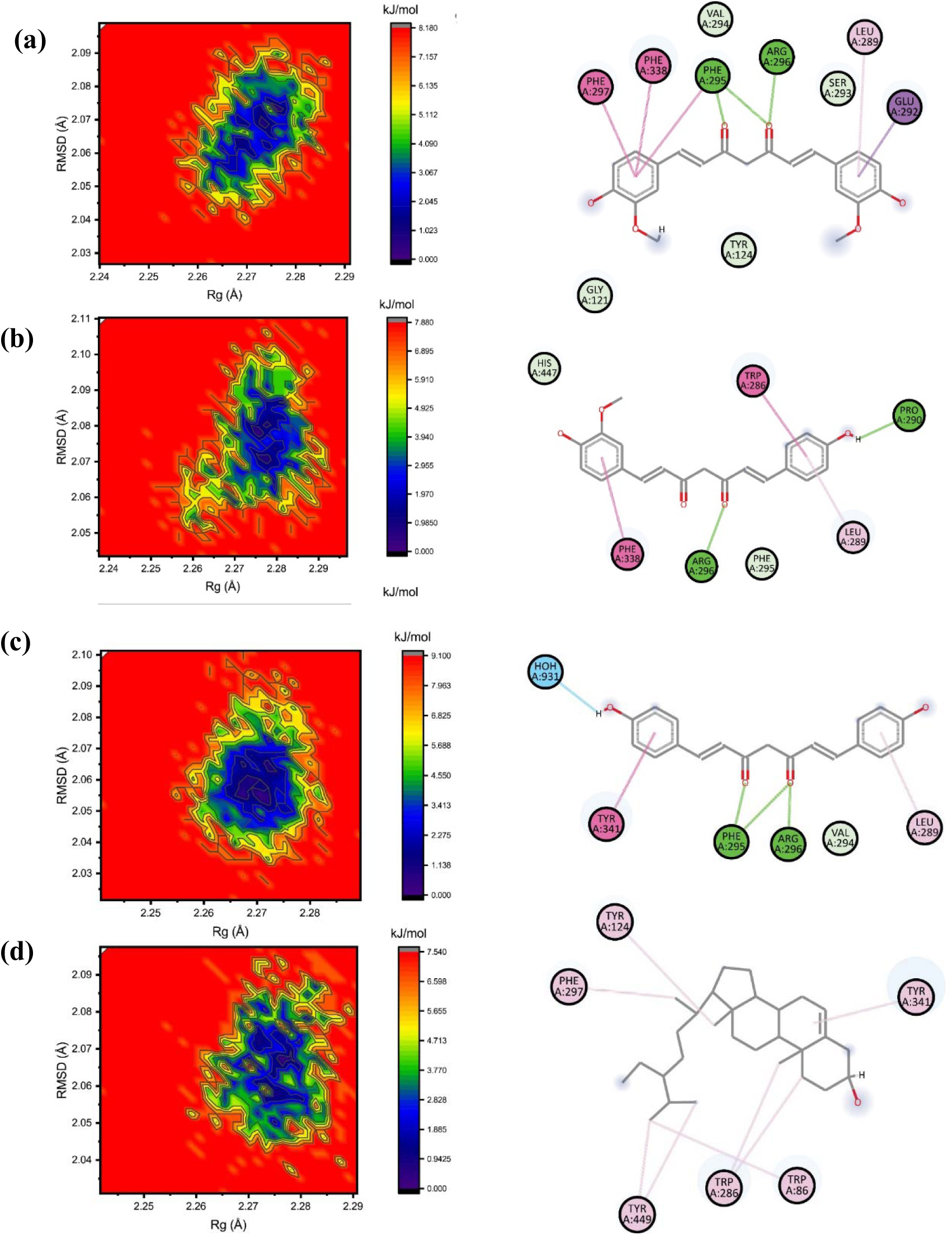
The Free Energy Landscape Plots of top four ligands (a) Curcumin, (b) Demethoxycurcumin, (c) Bis Demethoxycurcumin and (d) Beta-sitosterol. Here. blue regions describe the conformations with lower energy while, the red regions describe the conformations with higher energy. The color bar represents the relative free-energy value in kcal mol^−1^.

## 4. Discussion

Use of natural compounds to design new drugs has some tremendous advantages over the chemically synthesized drugs as they have minimal side effects, unique chemical structures, less long term toxicity, better bioavailability and biological activities. Natural products are being used from the Vedic periods. It is also true that about 80% of available drugs in the market are inspired from natural products (72). In western pharmaceutical company natural products got much attention during the period of 1970-1980 (73) and almost half of the drugs approved since 1981 are derived from natural products (74). *C. amada* has been used from the ancient period for biliousness, asthma, skin diseases, itching and inflammation (30). Besides, several research demonstrated that *C. amada* compounds have ability to fight against mouth and ear inflammation, ulcers, and stomatitis (75, 76). Molecular docking models the interaction between the ligand and receptor protein at their atomic levels and also provides us the information about the conformation and behavior of small molecules inside the macromolecules (77). Chemical profiling provides us the details information about the indigenous chemical compounds of any source and their detailed properties. The significant chemical compounds can be screened out from thousands of compounds by computer algorithms termed as virtual screening (78). In today’s drug discovery process virtual screening got much attention compared to the empirical screening due to the robustness, cost effective and time saving nature of this method (78). The virtual screening and precision docking of four ligands predicted the better binding affinity of curcumin towards AChE than demethoxycurcumin, bisdemethoxycurcumin and β-sitosterol. According to the docking interactions, curcumin formed two hydrogen bonds with the selective PHE^295^ residue, thus can inhibit the access of choline ester series. Promisingly, curcumin was able to form pi-alkyl bond with TYR^337^ residue of PAS site which is required for ligand recognition. In the same manner demethoxycurcumin, bisdemethoxy curcumin formed two hydrogen bonds with PHE^295^ residue. But, In comparison with the above compounds, beta-sitosterol did not form any hydrogen bond but hydrophobic interactions with PAS site, selective and conserved residues. This compound formed three pi-alkyl interactions with PAS site residue TRP^86^ and two pi-alkyl bonds with selective TRP^286^ residue. As mentioned above, the PAS site residues TRP^86^, TYR^133^, TYR^337^ and PHE^338^ have significant role in ligand identification and allosteric activation (79, 80). The stability of the substrate in this site is achieved by pi-cation interaction whereas, the selectivity of the substrates is achieved by preventing the access of choline ester series, medicated by PHE^295^ and PHE^297^ (81). A comprehensive analysis of enzyme inhibitor complexes has shown that the indole ring of TRP^286^ interacts directly with several inhibitors with a variety of modes of interaction that depend on the nature of ligands, including stacking, aromas, and p-cation (82, 83). Molecular dynamics simulation mimics what atoms do in real biological system and it is an effective way to understand the macromolecular structure to function relationship (84). MDS provides information about the interaction, stability of ligand inside the receptor proteins and conformational changes of ligand or macromolecules that may undergo inside the biological system in various conditions with the course of time (85). The structural analysis through MD Simulation also followed the Molecular docking prediction. In MD simulation, hydrogen bond is a major factor by which the stability of molecules. Curcumin showed the maximum hydrogen bond in complexed with AChE than other three ligands so that curcumin appeared the stable conformation than demethoxycurcumin, bisdemethoxycurcumin and β-sitosterol as the higher the hydrogen bond, the stable the molecules. As the free energy conducts the molecular processes inside the biological system thus, binding free energy determination is one of the most important task in molecular studies. The MM-PBSA methods is widely used due to its robust and modular nature (86). Though the β-sitosterol showed the highest binding energy with more fluctuations during the simulation, it could not be the best inhibitor against AChE because all other results such as RMSD, RMSF displayed higher values due to lower hydrogen bonding. Many new developed drugs failed to enter into the market due to their low pharmacokinetic activity (87). Therefore, identifying lead compounds with better absorption properties, potent site of action without side effects are most preferable. The compounds bioavailability, drug likeness and toxicity were predicted by means of ADMET (absorption, distribution, metabolism, elimination and toxicity) (88, 89) parameters. Due to low cost and time saving properties of computer-based ADMET technique compared to the standard laboratory experiment gained much attention to the scientists (49, 52). New leads or drug compounds prediction for CNS diseases must possess some vital physicochemical factors. These vital parameters acceptable values are depicted in table S-3 (56, 90). Molecular weight is an important factor to screen out probable drug compounds because higher molecular weight containing compounds can’t pass the blood brain barrier and also they have lower drug likeness activity (50). In our study we applied the drug likeness of the selected compounds was evaluated by Lipinski’s “Rule of Five”, ro5 (50). According this law our selected four compounds showed no violations of these rules, but one compound β-Sitosterol violate one parameter of “ro5” (shown in supplementary table 4). The Molecular weight, HBD and HBA of these compounds were in the acceptable range. Above the standards range of these parameters the compounds have worse permeability. All of the compounds aquatic solubility and caco cell permeability were in acceptable range instead of compound β-Sitosterol. β-Sitosterol aqueous solubility was below the acceptable range. The HOA percentage result depicts that’s our selected compounds (Bisdemethoxycurcumin, Curcumin Demethoxycurcumin and β-Sitosterol have the following higher permeability 81.57%, 87%, 79.57% and 100% respectively (supplementary table 4). The log B/B predicted result showed that all of our selected compounds were in the acceptable range (supplementary table 4). This assessment is necessary because higher polar compound can’t pass BBB (51, 52). The compounds distribution was simulated by human serum albumin binding affinities calculation (HSA). This calculation showed that a significant proportion of compounds can circulate freely through the blood stream and thus reach the drug target sites. Except, B-sitosterol all of our selected three compounds have this ability to move freely in the blood stream. This curcuma derived compounds can be used as a good AChE inhibitor for cognitive impairment treatment and could open a new door for NDDs treatment. Further, in future, potential inhibitor design research based on the improved and optimized structural similarity of curcumin, bisdemethoxycurcumin and demethoxycurcumin could be better option for NDDs treatment line.

## 5. Conclusion

Neurodegenerative diseases are posing severe threats to human existence. The market available drugs have some serious side effects. So, search of new natural compounds which can fight against NDD with minimum side effects are mostly preferable. Hence, in our study Curcuma derived compounds are studied to inhibit AChE as the AChE monomers prolong the formation of Aβplaques and responsible for cognitive impairment. The MD simulation explained that the Curcumin was the most favorable inhibitor with higher binding free energy among the curcuma compounds. It showed the strongest configuration by binding interactions with the active site residue PHE 295 which is important for drug binding and activity. In addition, this study predicts that Curcumin, Bisdemethoxycurcumin and demethoxycurcumin was observed to hold potential for hERG toxicity while, Beta-sitosterol did not possess potential toxicity for hERG according to ADMET result. Thus, ADMET/T anticipated the curcumin and Bisdemethoxycurcumin were not toxic and can be more stable while binding with AChE compared to ligand Beta-sitosterol and finally, inhibition of AChE by Curcumin as well as Bisdemethoxycurcumin ligands will play an important role in the treatment of NDD diseases.

## Supporting information

Compounds structure

## Author Contributions

Raju Dash conceived the idea and designed the experimental work. Md. Chayan Ali and Yeasmin Akter Munni carried out the experiments. Md. Chayan Ali and Yeasmin Akter Munni, Raju Das, Marium sultana, Nasrin Akter, Mahbubur Rahman, Md Nazim Uddin, Kantu Das, Md. Hossen collected the data. Md. Chayan Ali and Yeasmin Akter Munni analyze the data and prepared the figures. Md. Chayan Ali and Yeasmin Akter Munni made the initial draft. All authors reviewed the final manuscript.

