## Supplementary material for "In silico chemical profiling and identification of neuromodulators from *Curcuma amada* targeting Acetylcholinesterase": Compounds structure

| **Compound Name** | **IUPAC Name** | **Structure** | **Binding Affinity Kcal/mol** |
| --- | --- | --- | --- |
| Bisdemethoxy Curcumin | (1E,6E)-1,7-bis(4-hydroxyphenyl)hepta-1,6-diene-3,5-dione | 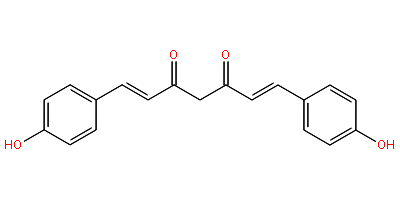 | -10.9 |
| beta-Sitosterol | (3S,8S,9S,10R,13R,14S,17R)-17-[(2R,5R)-5-ethyl-6-methylheptan-2-yl]-10,13-dimethyl-2,3,4,7,8,9,11,12,14,15,16,17-dodecahydro-1H-cyclopenta[a]phenanthren-3-ol | 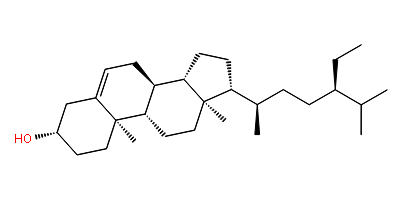 | -10 |
| alpha-Selinene | (3R,4aR,8aR)-5,8a-dimethyl-3-prop-1-en-2-yl-2,3,4,4a,7,8-hexahydro-1H-naphthalene | 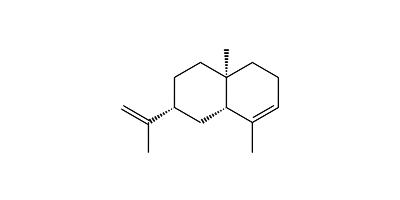 | -9.5 |
| Guaia-6,9-diene | (1S,3aR,8aR)-1,4-dimethyl-7-propan-2-yl-1,2,3,3a,6,8a-hexahydroazulene | 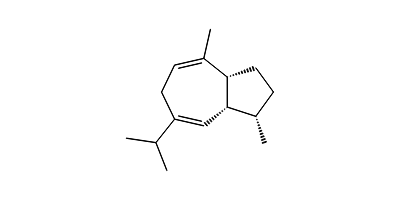 | -9.1 |
| alpha-Mururolene | (1S,4aS,8aR)-4,7-dimethyl-1-propan-2-yl-1,2,4a,5,6,8a-hexahydronaphthalene | 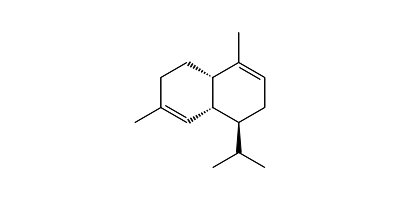 | -8.8 |
| Patchoulane | (3R,3aS,5R,8R,8aS)-3,8,9,9-tetramethyloctahydro-1H-5,8a-methanoazulene | 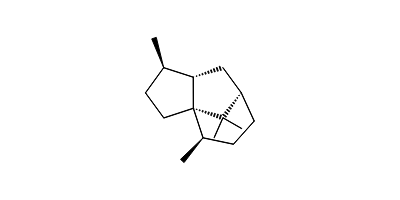 | -8.8 |
| beta-Copaene | (1S,6S,7S,8S)-8-isopropyl-1-methyl-3-methylenetricyclo[4.4.0.02,7]decane | 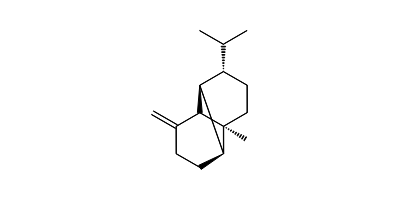 | -8.7 |
| Ledol | (4S,7S)-1,1,4,7-tetramethyl-2,3,4a,5,6,7,7a,7b-octahydro-1aH-cyclopropa[e]azulen-4-ol | 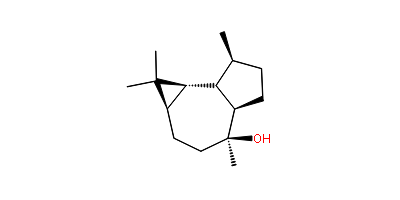 | -8.7 |
| (E)-Sabinol | (1S,3R)-4-methylidene-1-propan-2-ylbicyclo[3.1.0]hexan-3-ol | 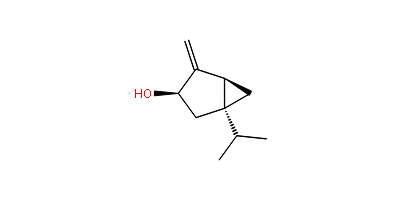 | -8.6 |
| 2-Hydroxy-2′-Methoxy diphenyl ether / 2-(2-methoxyphenoxy)phenol | 2-(2-methoxyphenoxy)phenol | 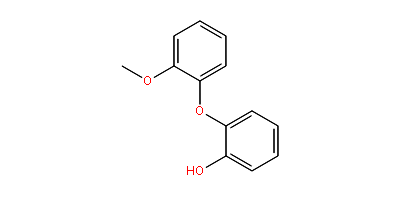 | -8.5 |
| delta-Elemene | (3R,4R)-4-ethenyl-4-methyl-1-propan-2-yl-3-prop-1-en-2-ylcyclohexene | 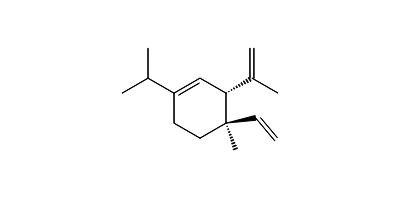 | -8.4 |
| Spathulenol | (1aR,4aR,7S,7aR,7bR)-1,1,7-trimethyl-4-methylidene-1a,2,3,4a,5,6,7a,7b-octahydrocyclopropa[h]azulen-7-ol | 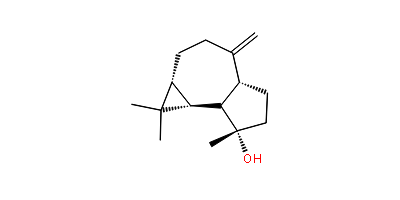 | -8.4 |
| (E)-beta-Farnesene | (6E)-7,11-dimethyl-3-methylidenedodeca-1,6,10-triene | 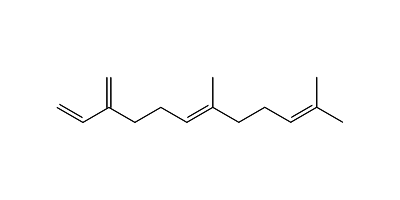 | -8.3 |
| Zerumin B | 4-[(1R)-2-[(1S,4aS,8aS)-5,5,8a-trimethyl-2-methylidene-3,4,4a,6,7,8-hexahydro-1H-naphthalen-1-yl]-1-hydroxyethyl]-2-hydroxy-2H-furan-5-one | 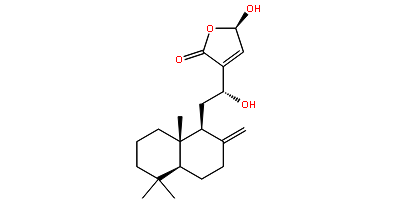 | -8.3 |
| Epicurzerenone | (5R,6S)-6-ethenyl-3,6-dimethyl-5-prop-1-en-2-yl-5,7-dihydro-1-benzofuran-4-one | 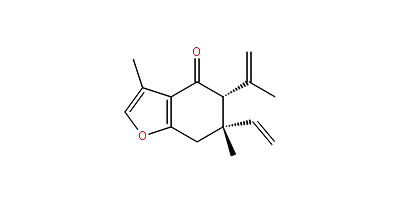 | -8.1 |
| cis-Farnesal | (2Z,6E)-3,7,11-trimethyldodeca-2,6,10-trienal | 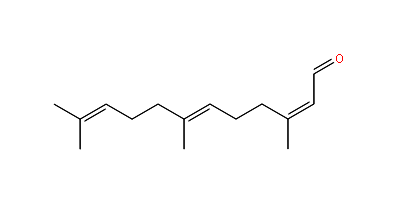 | -8.1 |
| Car-3-ene | (1S,6R)-4,7,7-trimethylbicyclo[4.1.0]hept-3-ene | 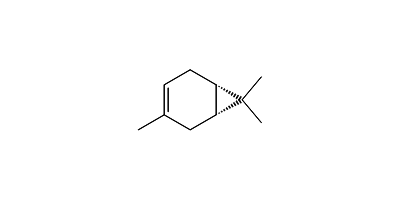 | -8 |
| alpha-Ionone | (E)-4-(2,6,6-trimethylcyclohex-2-en-1-yl)but-3-en-2-one | 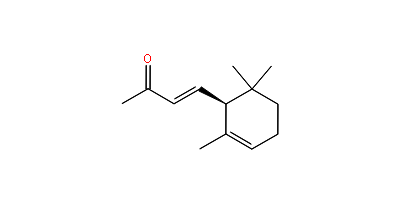 | -7.9 |
| (Z)-beta-Farnesene | (6Z)-7,11-dimethyl-3-methylidenedodeca-1,6,10-triene | 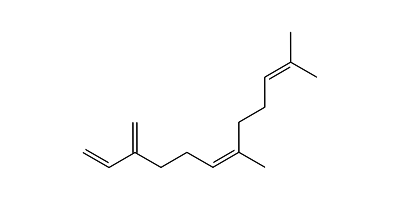 | -7.9 |
| gamma-Elemene | (1S,2S)-1-ethenyl-1-methyl-4-propan-2-ylidene-2-prop-1-en-2-ylcyclohexane | 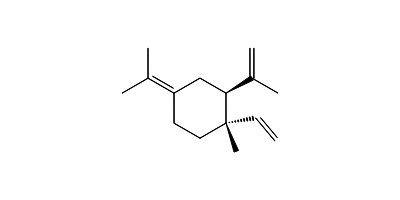 | -7.8 |
| alpha-Terpinyl acetate | 2-(4-methylcyclohex-3-en-1-yl)propan-2-yl acetate | 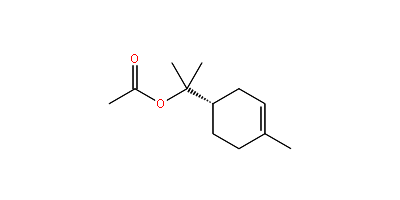 | -7.7 |
| Zerumbone | (2E,6E,10E)-2,6,9,9-tetramethylcycloundeca-2,6,10-trien-1-one | 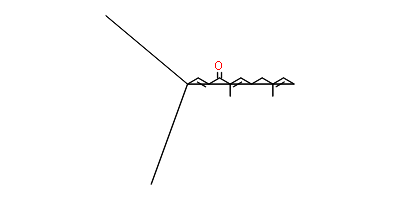 | -7.7 |
| Caffeic acid | (E)-3-(3,4-dihydroxyphenyl)prop-2-enoic acid | 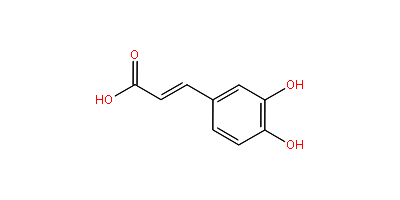 | -7.6 |
| alpha-Humulene | (1E,4E,8E)-2,6,6,9-tetramethylcycloundeca-1,4,8-triene | 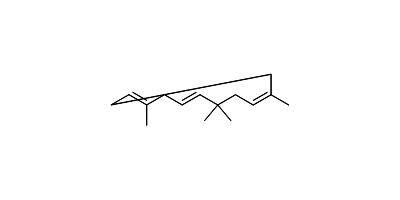 | -7.5 |
| Amadaldehyde | (17E,19E)-54-ethoxytrihexaconta-17,19-dien-7,9-diynal | 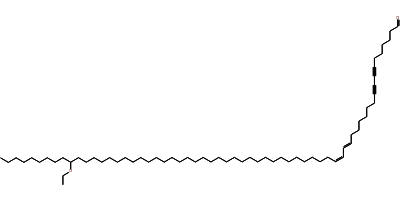 | -7.5 |
| Labdadiene | (2E)-2-[2-(5,5,8a-trimethyl-2-methylidene-3,4,4a,6,7,8-hexahydro-1H-naphthalen-1 yl)ethylidene]butanedial | 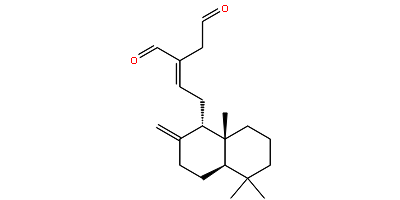 | -7.4 |
| p-Coumaric acid | (E)-3-(4-hydroxyphenyl)prop-2-enoic acid | 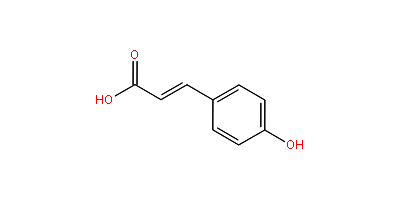 | -7.4 |
| alpha-Phellandrene | 2-methyl-5-propan-2-ylcyclohexa-1,3-diene | 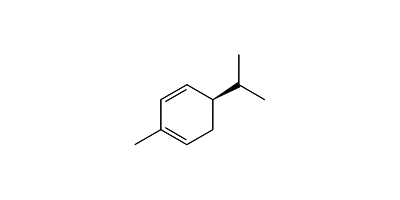 | -7.3 |
| Terpinolene | 1-methyl-4-propan-2-ylidenecyclohexene | 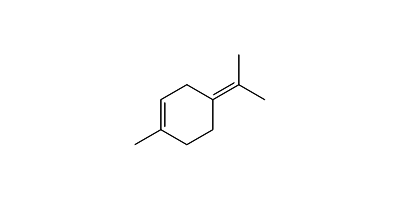 | -7.3 |
| Camphene | 3,3-dimethyl-2-methylidenebicyclo[2.2.1]heptane | 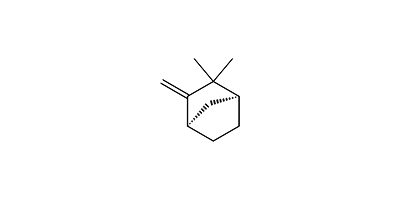 | -7.2 |
| alpha-Terpinene | 1-methyl-4-propan-2-ylcyclohexa-1,3-diene | 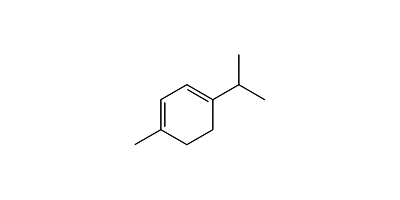 | **-7.2** |
| gamma-Terpinene | 1-methyl-4-propan-2-ylcyclohexa-1,4-diene | 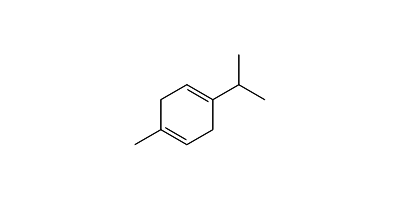 | **-7.2** |
| Pinocamphone | 4,6,6-trimethylbicyclo[3.1.1]heptan-3-one | 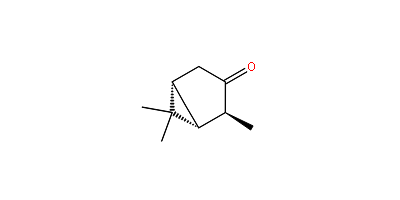 | **-7.2** |
| trans-Pinocarveol | (1R,3S,5R)-6,6-dimethyl-4-methylidenebicyclo[3.1.1]heptan-3-ol | 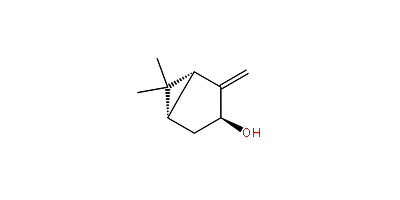 | **-7.2** |
| Carvomenthene | 1-methyl-4-propan-2-ylcyclohexene | 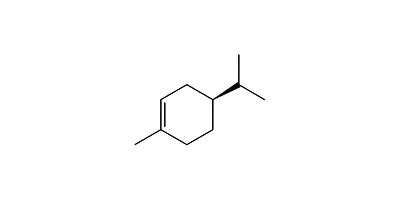 | **-7.1** |
| beta-Phellandrene | 3-methylidene-6-propan-2-ylcyclohexene | 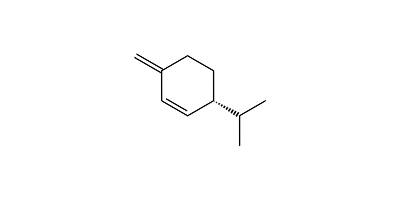 | **-7.1** |
| alpha-Terpineol | 2-(4-methylcyclohex-3-en-1-yl)propan-2-ol | 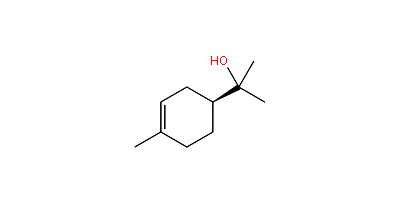 | -7.1 |
| beta-Elemene | (1S,2S,4R)-1-ethenyl-1-methyl-2,4-bis(prop-1-en-2-yl)cyclohexane | 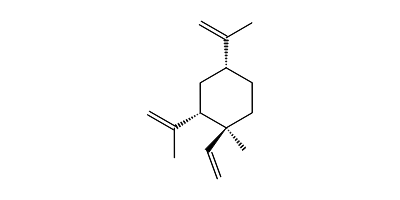 | -7.1 |
| Cinnamic acid | (E)-3-phenylprop-2-enoic acid | 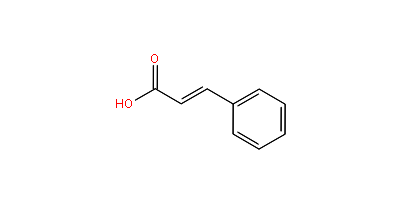 | -7.1 |
| Limonene | 1-methyl-4-prop-1-en-2-ylcyclohexene | 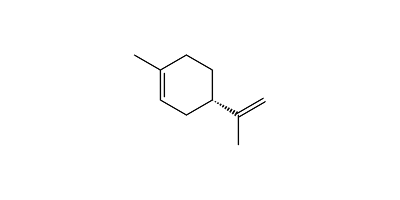 | -7 |
| Octadecane | octadecane | 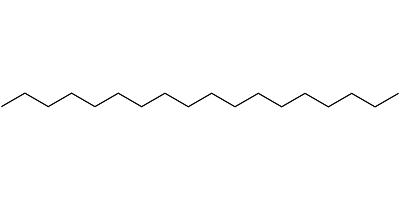 | -7 |
| alpha-Bergamotene | 4,6-dimethyl-6-(4-methylpent-3-enyl)bicyclo[3.1.1]hept-3-ene | 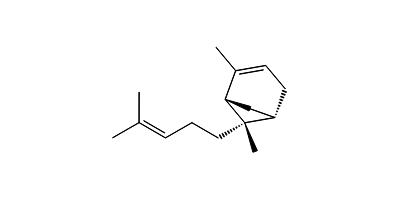 | -7 |
| Perillyl alcohol | (4-prop-1-en-2-ylcyclohexen-1-yl)methanol | 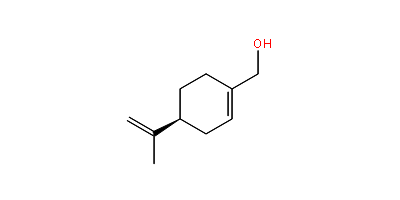 | -7 |
| beta-(E)-Caryophyllene | (1R,9S,E)-4,11,11-trimethyl-8-methylenebicyclo[7.2.0]undec-4-ene | 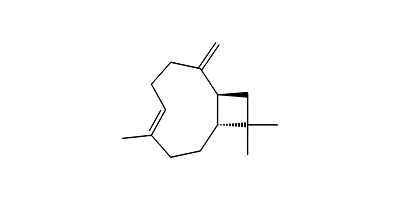 | -7 |
| Acetone | propan-2-one | 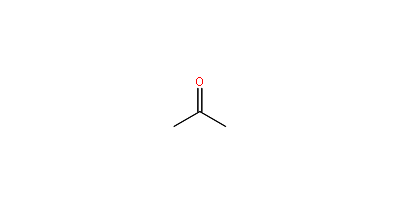 | -6.9 |
| p-Menth-1-en-9-ol | 2-(4-methylcyclohex-3-en-1-yl)propan-1-ol | 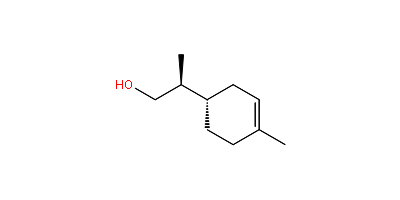 | -6.9 |
| beta-Terpineol | 1-methyl-4-prop-1-en-2-ylcyclohexan-1-ol | 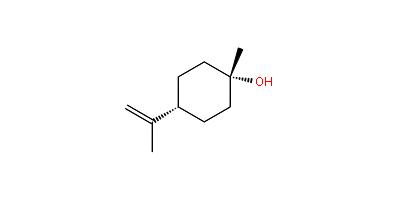 | -6.9 |
| Geranyl formate | [(2E)-3,7-dimethylocta-2,6-dienyl] formate | 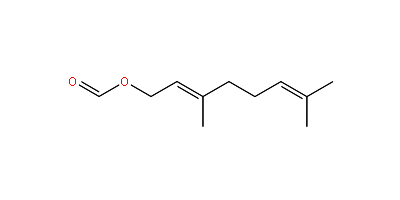 | -6.9 |
| cis-Ocimene | (3Z)-3,7-dimethylocta-1,3,6-triene | 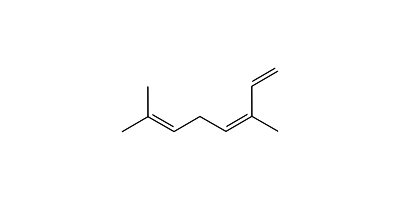 | -6.8 |
| Ocimene | (3E)-3,7-dimethylocta-1,3,6-triene | 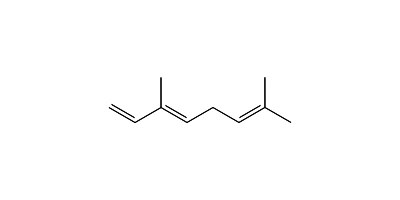 | -6.8 |
| p-Menth-4-en-9-ol | 2-(4-methylcyclohexen-1-yl)propan-1-ol | 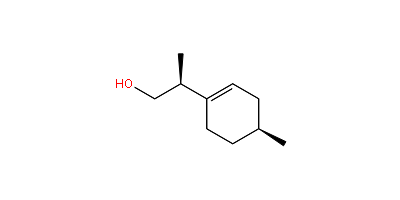 | -6.8 |
| Citral | (2E)-3,7-dimethylocta-2,6-dienal | 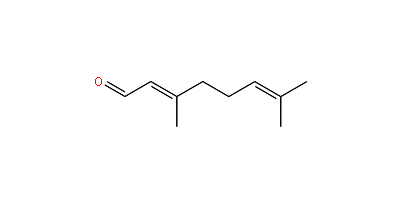 | -6.8 |
| trans-Sabinene hydrate | (1R,4S)-4-methyl-1-propan-2-ylbicyclo[3.1.0]hexan-4-ol | 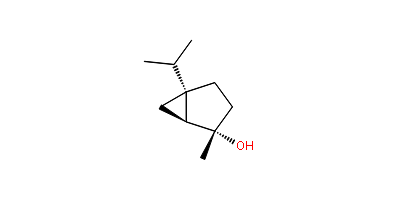 | -6.8 |
| alpha-Fenchol | (1R,3S,4S)-2,2,4-trimethylbicyclo[2.2.1]heptan-3-ol | 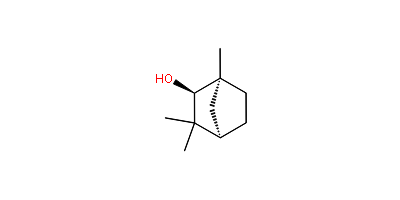 | -6.8 |
| Vanillic acid | 4-hydroxy-3-methoxybenzoic acid | 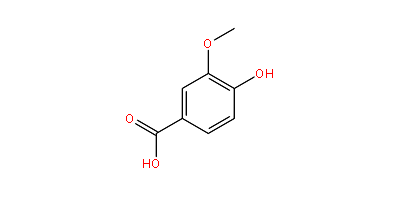 | -6.8 |
| Cubebol | (3S,3aR,3bR,4S,7R,7aR)-4-isopropyl-3,7-dimethyloctahydro-1H-cyclopenta[1,3]cyclopropa[1,2]benzen-3-ol | 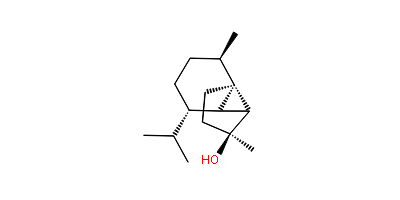 | -6.8 |
| Myrtenol | (6,6-dimethyl-4-bicyclo[3.1.1]hept-3-enyl)methanol | 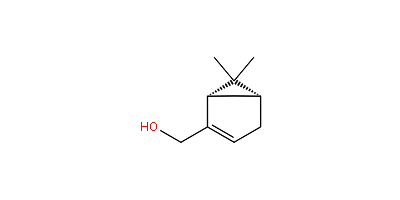 | -6.7 |
| Sabinene | 4-methylidene-1-propan-2-ylbicyclo[3.1.0]hexane | 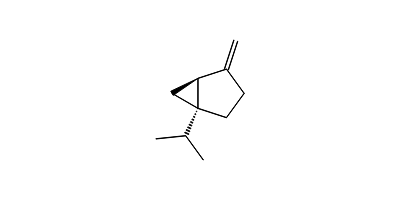 | -6.6 |
| 2-Methyl-6-methyleneocta-1,7-dien-3-one / 2,6-Dimethyleneoct-7-en-3-one | 2-methyl-6-methylideneocta-1,7-dien-3-one | 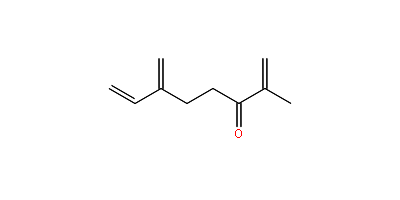 | -6.6 |
| Norbornyl acetate | 3-bicyclo[2.2.1]heptanyl acetate | 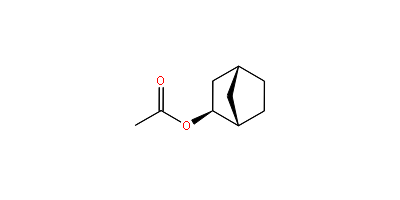 | -6.6 |
| Lavandulol | (2R)-5-methyl-2-prop-1-en-2-ylhex-4-en-1-ol | 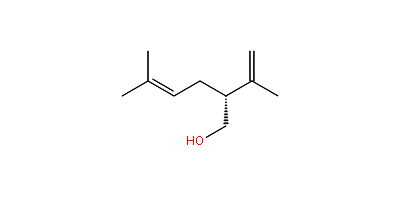 | -6.6 |
| Borneol | 4,7,7-trimethylbicyclo[2.2.1]heptan-3-ol | 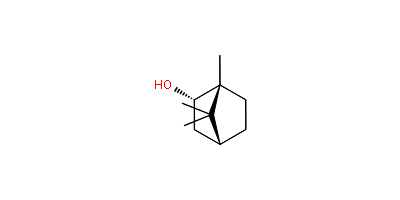 | -6.6 |
| 2-Methyl-6-methylene-3,7-octadien-2-ol | 2-methyl-6-methyleneocta-3,7-dien-2-ol | 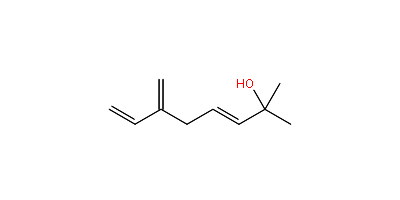 | -6.6 |
| Gallic acid | 3,4,5-trihydroxybenzoic acid | 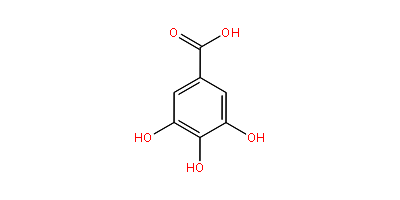 | -6.6 |
| Syringic acid | 4-hydroxy-3,5-dimethoxybenzoic acid | 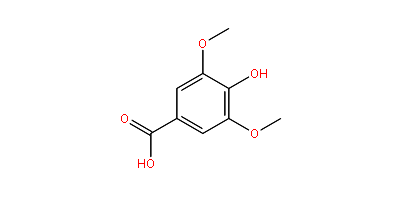 | -6.6 |
| beta-Pinene | 6,6-dimethyl-4-methylidenebicyclo[3.1.1]heptane | 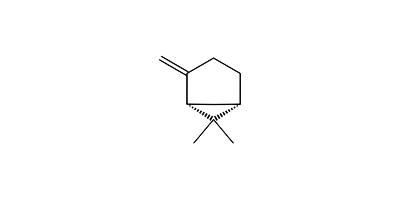 | -6.5 |
| trans-Dihydroocimene | (3E)-3,7-dimethylocta-1,3-diene | 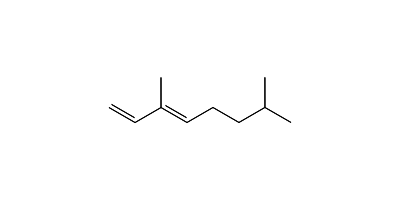 | -6.5 |
| Linalool | 3,7-dimethylocta-1,6-dien-3-ol | 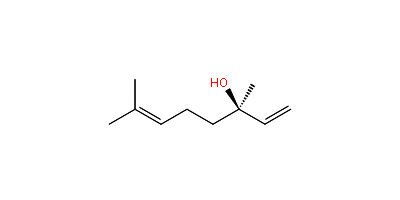 | -6.5 |
| Hexadecane | hexadecane | 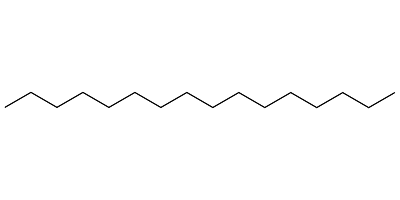 | -6.4 |
| Myrcenol | 2-methyl-6-methylideneoct-7-en-2-ol | 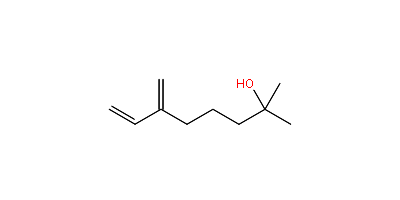 | -6.4 |
| 2,6-Dimethylhept-5-en-1-al | 2,6-dimethylhept-5-enal | 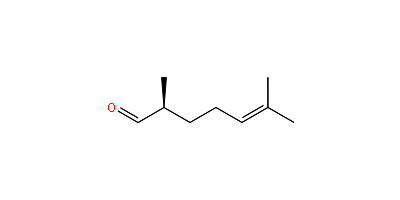 | -6.1 |
| (Ethoxymethyl)benzene | ethoxymethylbenzene | 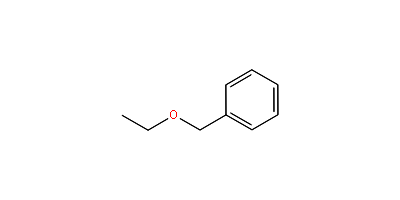 | -6.1 |
| Hept-1-en-1-ol acetate | (E)-hept-1-en-1-yl acetate | 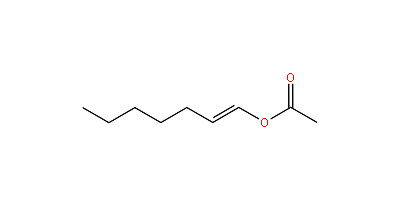 | -6.1 |
| Undecan-2-one | undecan-2-one | 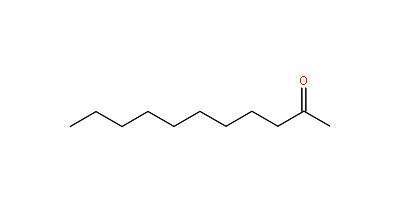 | -6.1 |
| Undecanal | undecanal | 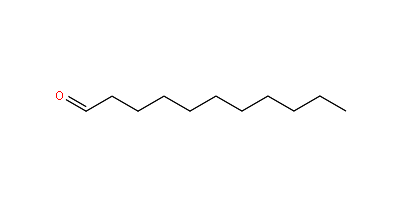 | -5.9 |
| Heptadecane | heptadecane | 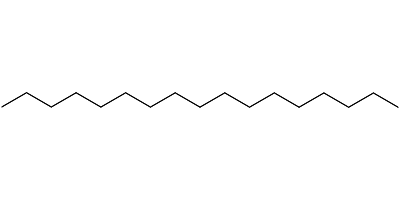 | -5.9 |
| Nonan-2-ol | nonan-2-ol | 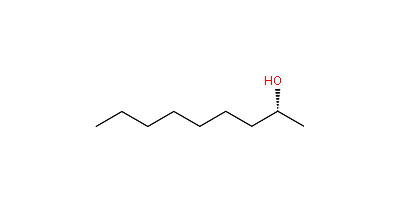 | -5.8 |
| 2-Methylheptan-3-ol | 2-methylheptan-3-ol | 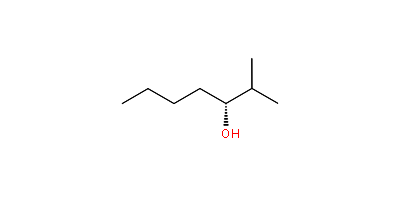 | -5.7 |
| Non-2-en-4-one | (E)-non-2-en-4-one | 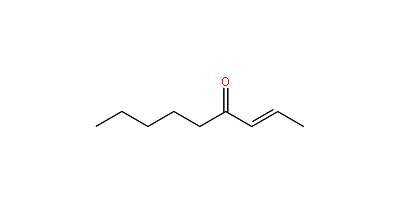 | -5.6 |
| alpha-Pinene | 4,6,6-trimethylbicyclo[3.1.1]hept-3-ene | 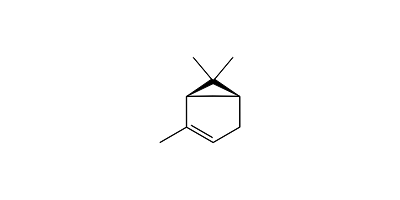 | -3.3 |
| Curcumin | (1E,6E)-1,7-bis(4-hydroxy-3-methoxyphenyl)hepta-1,6-diene-3,5-dione | 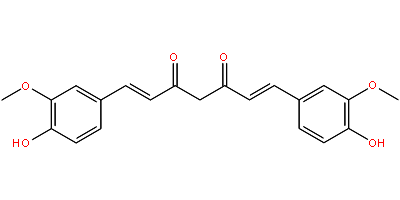 | -10.8 |
| beta-Gurjunene | (1aR,4R,4aR,7aR,7bR)-1,1,4-trimethyl-7-methylidene-2,3,4,4a,5,6,7a,7b-octahydro-1aH-cyclopropa[e]azulene | 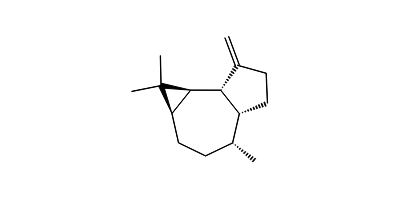 | -9.8 |
| Demethoxy Curcumin | (1E,6E)-1-(4-hydroxy-3-methoxyphenyl)-7-(4-hydroxyphenyl)hepta-1,6-diene-3,5-dione | 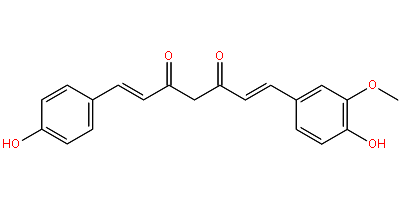 | -10.6 |
| (E)-Labda-8(17),13-diene-15,16-olide | 3-[2-[(1S,8aS)-5,5,8a-trimethyl-2-methylidene-3,4,4a,6,7,8-hexahydro-1H-naphthalen-1-yl]ethyl]-2H-furan-5-one | 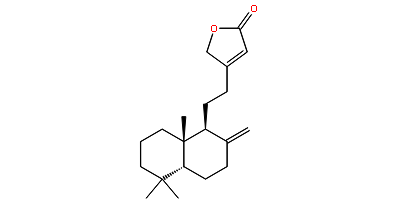 | -9.8 |
| beta-Selinene | (3R,4aS,8aR)-8a-methyl-5-methylidene-3-prop-1-en-2-yl-1,2,3,4,4a,6,7,8-octahydronaphthalene | 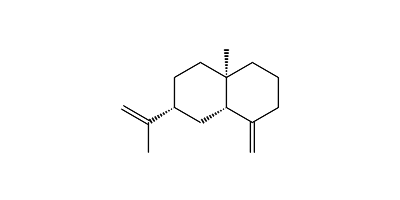 | -9.2 |
| beta-Curcumene | 1-methyl-4-[(2R)-6-methylhept-5-en-2-yl]cyclohexa-1,4-diene | 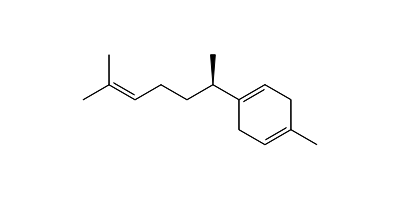 | -9.1 |
| ar-Turmerone / Turmerone | (6S)-2-methyl-6-(4-methylphenyl)hept-2-en-4-one | 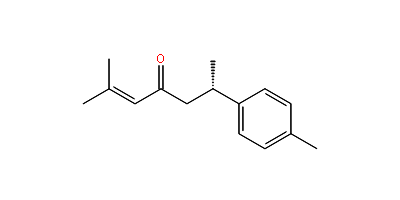 | -9 |
| beta-Bisabolol | 4-methyl-1-(6-methylhept-5-en-2-yl)cyclohex-3-en-1-ol | 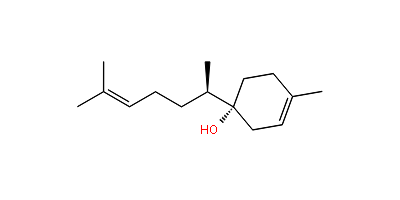 | -9 |
| Amadannulen | ethyl 3-(2-hydroxy-10-methyl-2,3,4,5,6,7,8,9,10,11-decahydro-1H-cyclopenta[10]annulen-5-yl)-5-methylcyclohexane-1-carboxylate | 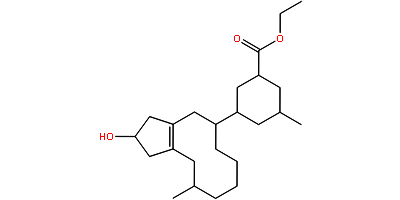 | -8.8 |
| alpha-Curcumene / ar-Curcumene | 1-methyl-4-(6-methylhept-5-en-2-yl)benzene | 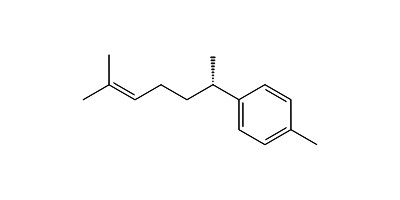 | -8.8 |
| Difurocumenonol | (11E)-13,15,23,25-tetrahydroxy-1,5,10,14,17,21-hexamethyl-7,19-dioxahexacyclo[13.9.1.02,14.04,8.016,24.018,22]pentacosa-4(8),5,11,16(24),18(22),20-hexaen-3-one | 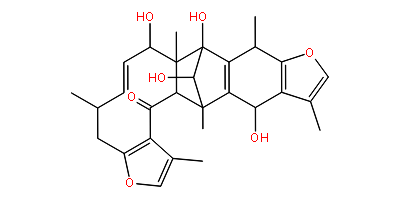 | -8.8 |
| alpha-Copaene | (1S,6S,7S,8S)-8-isopropyl-1,3-dimethyltricyclo[4.4.0.02,7]dec-3-ene | 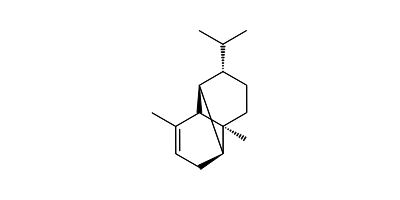 | -8.7 |
| gamma-Guaiene | (3R,8R)-5-isopropyl-3,8-dimethyl-1,2,3,6,7,8-hexahydroazulene | 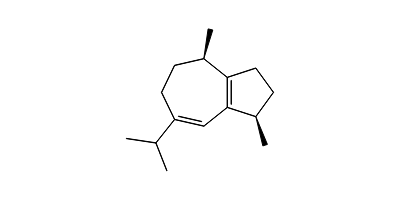 | -8.7 |
| Coronarin B | 7-[(1S,4aS,8aS)-5,5,8a-trimethyl-2-methylidene-3,4,4a,6,7,8-hexahydro-1H-naphthalen-1-yl]-3-hydroxy-4,7-dihydro-3H-dioxepine-5-carbaldehyde | 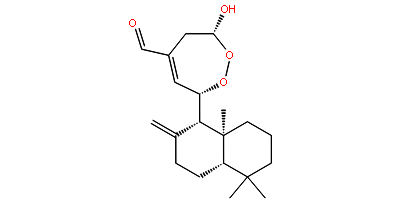 | -8.7 |
| (E,E)-alpha-Farnesene | "(3E,6E)-3,7,11-trimethyldodeca-1,3,6,10-tetraene | 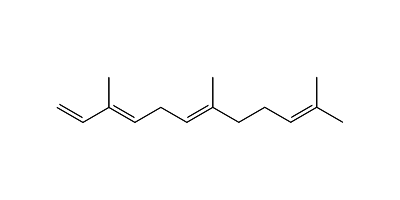 | -8.6 |
| Zederone | (1aR,10aR,Z)-3,7,10a-trimethyl-6,9,10,10a-tetrahydrooxireno[2',3':4,5]cyclodeca[1,2-b]furan-2(1aH)-one | 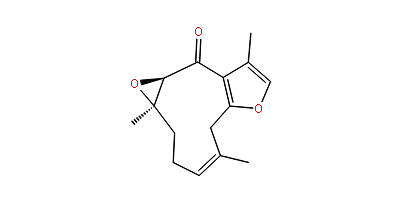 | -8.6 |
| Caryophyllene | "(1R,4E,9S)-4,11,11-trimethyl-8-methylidenebicyclo[7.2.0]undec-4-ene | 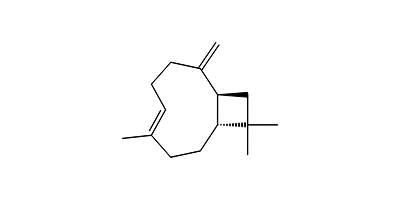 | -8.5 |
| Curzerenone | "(5R,6R)-6-ethenyl-3,6-dimethyl-5-prop-1-en-2-yl-5,7-dihydro-1-benzofuran-4-one | 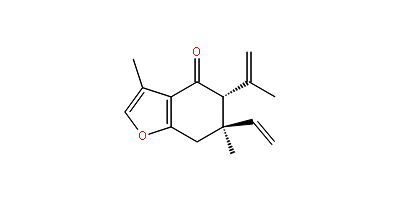 | -8.5 |
| alpha-Longipinene | (1R,2S,7R,8R)-2,6,6,9-tetramethyltricyclo[5.4.0.02,8]undec-9-ene | 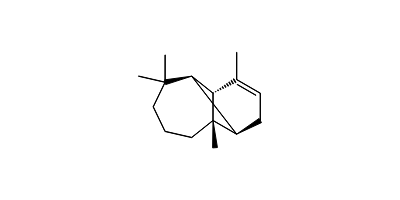 | -8.5 |
| Coronarin D | (3E)-3-[2-[(4aS,8aS)-5,5,8a-trimethyl-2-methylidene-3,4,4a,6,7,8-hexahydro-1H-naphthalen-1-yl]ethylidene]-5-hydroxyoxolan-2-one | 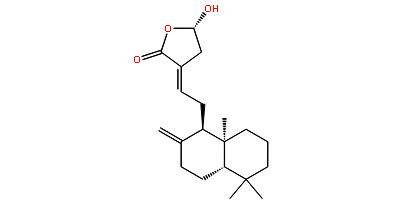 | -8.5 |
| alpha-Zingiberene / Zingiberene | (5R)-2-methyl-5-[(2S)-6-methylhept-5-en-2-yl]cyclohexa-1,3-diene |  | -8.4 |
| Germacrone | (3E,7E)-3,7-dimethyl-10-propan-2-ylidenecyclodeca-3,7-dien-1-one |  | -8.4 |
| p-Menth-4-en-9-ol | 2-(4-methylcyclohexen-1-yl)propan-1-ol |  | -8.4 |
| trans-Farnesal | (2E,6E)-3,7,11-trimethyldodeca-2,6,10-trienal |  | -8.3 |
| Bisabolene | (4Z)-1-methyl-4-(6-methylhept-5-en-2-ylidene)cyclohexene |  | -8.1 |
| Curzerene | (5R,6R)-6-ethenyl-3,6-dimethyl-5-prop-1-en-2-yl-5,7-dihydro-4H-1-benzofuran |  | -8.1 |
| alpha-guaiene | (1S,4S,7R)-1,4-dimethyl-7-prop-1-en-2-yl-1,2,3,4,5,6,7,8-octahydroazulene |  | -7.9 |
| Zerumin A | (Z)-5-[(1S,4aS,8aS)-5,5,8a-trimethyl-2-methylidene-3,4,4a,6,7,8-hexahydro-1H-naphthalen-1-yl]-3-formylpent-3-enoic acid |  | -7.8 |
| Ferulic acid | (E)-3-(4-hydroxy-3-methoxyphenyl)prop-2-enoic acid |  | -7.7 |
| delta-Car-3-ene / 3-Carene | 4,7,7-trimethylbicyclo[4.1.0]hept-3-ene |  | -7.5 |
| Thymol | 5-methyl-2-propan-2-ylphenol |  | -7.4 |
| p-Cymene | 1-methyl-4-propan-2-ylbenzene |  | -7.3 |
| alpha-Muurolol | (1R,4R,4aR,6S,8aR)-1,6-dimethyl-4-propan-2-yl-3,4,4a,5,6,7,8,8a-octahydro-2H-naphthalen-1-ol |  | -7.2 |
| Bornyl formate | (4,7,7-trimethyl-3-bicyclo[2.2.1]heptanyl) formate |  | -7.2 |
| alpha-Terpinene | 1-methyl-4-propan-2-ylcyclohexa-1,3-diene |  | -7.2 |
| p-Cymen-8-ol | 2-(4-methylphenyl)propan-2-ol |  | -7.1 |
| Caryophyllene oxide | (1R,4R,6R,10S)-4,12,12-trimethyl-9-methylene-5-oxatricyclo[8.2.0.04,6]dodecane |  | -7.1 |
| Camphor | 4,7,7-trimethylbicyclo[2.2.1]heptan-3-one |  | -7 |
| Geraniol | (2E)-3,7-dimethylocta-2,6-dien-1-ol |  | -7 |
| Myrcene | 7-methyl-3-methylideneocta-1,6-diene |  | -6.9 |
| p-Menth-1,8-dien-9-ol | 2-(4-methylcyclohex-3-en-1-yl)prop-2-en-1-ol |  | -6.9 |
| Gentisic acid | 2,5-dihydroxybenzoic acid |  | -6.9 |
| cis-Ocimene | (3Z)-3,7-dimethylocta-1,3,6-triene |  | -6.8 |
| Germacrene D | (1Z,6Z)-1-methyl-5-methylidene-8-propan-2-ylcyclodeca-1,6-diene |  | -6.8 |
| Myrtenal | 6,6-dimethylbicyclo[3.1.1]hept-3-ene-4-carbaldehyde |  | -6.8 |
| Neral | (2Z)-3,7-dimethylocta-2,6-dienal |  | -6.7 |
| Sabinene | 4-methylidene-1-propan-2-ylbicyclo[3.1.0]hexane |  | -6.6 |
| Terpinen-4-ol | "4-methyl-1-propan-2-ylcyclohex-3-en-1-ol |  | -6.6 |
| Citronellal | 3,7-dimethyloct-6-enal |  | -6.6 |
| (Z,Z)-Alloocimene | (4Z,6Z)-2,6-dimethylocta-2,4,6-triene |  | -6.6 |
| Protocatechuic acid | 3,4-dihydroxybenzoic acid |  | -6.6 |
| beta-Pinene | 6,6-dimethyl-4-methylidenebicyclo[3.1.1]heptane |  | -6.5 |
| Eucalyptol | 2,2,4-trimethyl-3-oxabicyclo[2.2.2]octane |  | -6.5 |
| Isoborneol | (1R,3R,4R)-4,7,7-trimethylbicyclo[2.2.1]heptan-3-ol |  | -6.5 |
| cis-Dihydroocimene | (3Z)-3,7-dimethylocta-1,3-diene |  | -6.3 |
| Dihydromyrcenol | 2,6-dimethyloct-7-en-2-ol |  | -6.1 |
| 6-Methyl-5-hepten-2-one | 6-methylhept-5-en-2-one |  | -6 |
| (E)-Thujone | (1S,4S,5R)-4-methyl-1-propan-2-ylbicyclo[3.1.0]hexan-3-one |  | -5.9 |
| Nonan-2-one / 2-Nonanone | nonan-2-one |  | -5.7 |
| alpha-Pinene | 4,6,6-trimethylbicyclo[3.1.1]hept-3-ene |  | -3.3 |
