## Supplementary table file for "In silico chemical profiling and identification of neuromodulators from *Curcuma amada* targeting Acetylcholinesterase"

**Table S1: Docking result of the selected compounds**

| **Molecule** | **Docking score** |
| --- | --- |
| Demethoxy curcumin | -11.913 |
| β-sitosterol | -10.729 |
| Bisdemethoxycurcumin | -12.997 |
| Curcumin | -14.523 |

**Table S2: Wet lab experimental effects of the selected four compounds**

| Compound | Conditions | References |
| --- | --- | --- |
| Curcumin | AD: safe and well tolerated; reduce skin inflammation, inhibits tumor formation | (1, 2) |
| Bisdemethoxycurcumin | Antiulcer effect, inducible nitric oxide synthase (iNOS) inhibition, antioxidant activity | (3, 4) |
| Demethoxycurcumin | PD: Antioxidative and Anti-inflammatory Properties in Parkinsonian Rats | (5) |
| Β-sitosterol | Modulates macrophage and rheumatoid inflammatory function in mice | (6) |

**Table S3: Vital ADMET parameters**

| **Attributes** | **Values** |
| --- | --- |
| Molecular weight (MW) | <500 |
| Hydrogen bond donors (HBD) | ≤ 5 |
| Hydrogen bond acceptors (HBA) | ≤ 10 |
| Predicted brain/blood partition coefficient (QPlog BB) | -3.0 to 1.2 |
| Predicted IC50 value for blockage of HERG K+ channels (QPlog herg) | < –5 |
| Human oral absorption (HOA) | 3=higher, 2= medium, 1=lower |
| Predicted apparent MDCK cell permeability in nm/sec (QPlog mdck) | <25 poor,  >500 great |

**Table S4: ADMET properties of selected compounds**

| ADMET properties | Molecules | | | |
| --- | --- | --- | --- | --- |
|  | Bisdemethoxycurcumin | Curcumin | Demethoxycurcumin | beta-Sitosterol |
| MW | 308.333 | 368.385 | 338.359 | 414.713 |
| HBD | 2 | 2 | 2 | 1 |
| HBA | 5.5 | 7 | 6.25 | 1.7 |
| QPlogPo/w | 2.577 | 2.922 | 2.523 | 7.472 |
| QPlogS | -4.028 | -4.439 | -3.996 | -8.314 |
| QPPCaco | 161.856 | 250.788 | 130.164 | 3429.643 |
| QPlogBB | -1.998 | -1.983 | -2.148 | -0.336 |
| HOA | 3 | 3 | 3 | 1 |
| SASA | 626.99 | 697.921 | 648.117 | 754.79 |
| FOSA | 85.495 | 263.477 | 151.384 | 676.384 |
| QPpolrz | 33.127 | 36.901 | 34.354 | 47.948 |
| QPlogPC16 | 12.261 | 12.985 | 12.539 | 12.473 |
| QPlogPoct | 16.924 | 18.647 | 17.821 | 17.64 |
| CIQPlogS | -3.996 | -4.616 | -4.308 | -7.033 |
| QPlogHERG | -6.44 | -6.286 | -6.134 | -4.53 |
| QPPMDCK | 69.106 | 110.936 | 54.603 | 1874.523 |
| QPlogKp | -2.788 | -2.533 | -3.069 | -1.638 |
| QPlogKhsa | -0.039 | -0.021 | -0.055 | 2.009 |
| #metab | 3 | 5 | 4 | 4 |
